# Rice Ethylene Receptors Function as Ca²⁺-Permeable Ion Channels to Orchestrate Calcium-Dependent Antagonism of Ethylene Responses in Roots

**DOI:** 10.1101/2025.10.20.683344

**Authors:** Zhangli Ye, Zijian Yang, Changyuan Li, Yangbo Chen, Enjie Yu, Chunhui Song, Zongran Yang, Shuo Liu, Hao Tian, Dongdong Kong, Legong Li, Liangyu Liu

## Abstract

The gaseous phytohormone ethylene governs essential plant processes spanning development, productivity, and stress resistance. As an essential nutrient and second messenger, calcium (Ca^2+^) is widely implicated in diverse plant physiological activities; however, its role in the ethylene signal transduction pathway remains elusive. Here, we identified a calcium-dependent antagonism of ethylene response (CAER) specifically modulates root elongation in the model cereal rice (*Oryza sativa*). Interestingly, we demonstrate that the ethylene receptors OsERS1 and OsERS2 both function as Ca^2+^-permeable channels, with OsERS1 further acting as a non-selective cation channel capable of permeating both monovalent and divalent cations. Mutagenesis analysis reveals that OsERS1 channel activity relies on homomeric assembly sites (C4/C6) rather than its ethylene-binding site (C65), indicating a clear decoupling of the molecular modules governing receptor signaling and ion channel function. The loss-of-function mutant *Osers1/2* does not exhibit the CAER phenotype of the wild type (WT), confirm that this calcium-dependent regulatory mechanism is dependent on both OsERS1 and OsERS2. These findings uncover an unexpected ion-channel function of ethylene receptors, redefining their molecular identity beyond canonical signaling proteins and establishing the novel “hormone receptor-type ion channel (HRIC)” paradigm—one that fundamentally expands our understanding of how plant hormones transduce signals at the membrane.

## INTRODUCTION

Phytohormones are small molecules that play crucial roles in regulating plant development and stress responses. Among them, the gaseous hormone ethylene (C_2_H_4_) regulates processes including cell division, seed germination, root elongation, senescence, flowering, and fruit ripening^1,2^. Ethylene also functions as a pivotal stress-responsive hormone, orchestrating plant adaptation to both biotic and abiotic stresses, including pathogen infection, drought, hypoxia, flooding, cold, heat and mechanical damage^3–8^. Canonically detected by membrane-localized receptors^8,9^, ethylene initiates signaling through a well-characterized linear pathway in plants: in the absence of ethylene, these receptors activate CONSTITUTIVE TRIPLE RESPONSE1 (CTR1), which in turn inhibits ETHYLENE INSENSITIVE2 (EIN2) and suppresses downstream signaling; upon ethylene binding, the hormone acts as an inverse agonist to attenuate receptor/CTR1 activity, thereby relieving EIN2 from inhibition and triggering transcriptional/translational reprogramming that underpins ethylene-mediated responses^10,11^.

Calcium (Ca^2+^) as a cellular second messenger is well-known in plant development and stress responses, and several studies have linked ethylene and Ca^2+^ signals^12,13^. Exogenous Ca^2+^ or internal calmodulin was found to suppress ethylene-mediated fruit ripening^14,15^, such as in papaya^16^, apple^17^, and litchi^18^. Similarly, ethylene-mediated flower senescence was also inhibited by calcium^19^. Notably, ethylene has been shown to activate uncharacterized membrane-localized calcium channels in tobacco protoplasts, leading to elevated cytosolic free calcium concentration ([Ca^2+^]_cyt_)^12^. However, the molecular identity of ethylene-regulated Ca^2+^ channels and the mechanistic integration of these pathways remain elusive.

Rice (*Oryza sativa*) is one of the top three staple crops worldwide in terms of yield, alongside wheat and maize. The ethylene-mediated “double response” in rice seedlings involves, characterized by inhibited primary root growth and promoted coleoptile elongation^20,21^, is a well-documented physiological phenomenon. Soil compaction elevates ethylene levels in rice roots, thereby impairing root penetration and elongation^22,23^; conversely, exogenous calcium application is agriculturally employed to promote rice germination and root elongation^24–26^. Notably, ethylene modulates root hair growth via fluctuations in [Ca^2+^]_cyt_ at root hair tips^27,28^. These observations underscore the clear physiological relevance of ethylene-Ca^2+^ crosstalk in rice, while highlighting the critical knowledge gap in its underlying mechanisms—particularly regarding how ethylene perception is mechanistically linked to Ca^2+^ signaling.

We found that the rice ethylene receptor OsERS1 (ETHYLENE RESPONSE SENSOR 1) possesses a novel function: it acts as a Ca^2+^-permeable cation channel, which significantly expands its functional scope beyond its role as an ethylene receptor. This finding establishes a close connection between ethylene and calcium signaling pathways and introduces a novel concept of “hormone receptor-type channel (HRIC)” in plants.

## RESULTS

### Crosstalk between calcium and ethylene signals

To investigate the interaction between Ca^2+^ and ethylene signals in rice, we first assessed the effect of Ca^2+^ on ethylene-mediated responses. Uniformly germinating seeds were placed on a wire mesh (1.6 cm above water with/without extra calcium chloride). The mesh was then put into an airtight container, where 2 ppm ethylene was injected via syringe, and liquid was sprayed to maintain humidity. The container was incubated in dark at 28°C for 3 days, with daily water replacement and ethylene replenishment^29^. Using etiolated hydroponic seedlings of the wild-type Dongjin (DJ) accession, we found that upon ethylene treatment in the absence of extra Ca^2+^ inhibited root growth by about 70% (Fig. 1a-c). However, when exogenous Ca^2+^ concentrations were increased from 1.5 mM to 5 mM, root elongation recovered to 60%-80% of the control level (Fig. 1a-c), demonstraing a phenomenon we term Calcium-dependent Antagonism of Ethylene Response (CAER) in rice roots. By contrast, no significant phenotypic change was observed in the coleoptile (Supplementary Fig. 1). These results indicate that Ca^2+^ helps rice roots counteract ethylene-induced inhibition, likely by interfering with ethylene signaling. Consistently, high Ca^2+^ suppressed the transcriptional upregulation of ethylene-activated marker genes such as OsERF073 (Fig. 1d and Supplementary Fig. 2). A similar CAER effect was observed in another rice accession, Nipponbare (Nip) (Supplementary Fig. 3). The CAER effect on root elongation in rice provides a useful system for studying calcium-ethylene signal interactions.

**Fig. 1.**
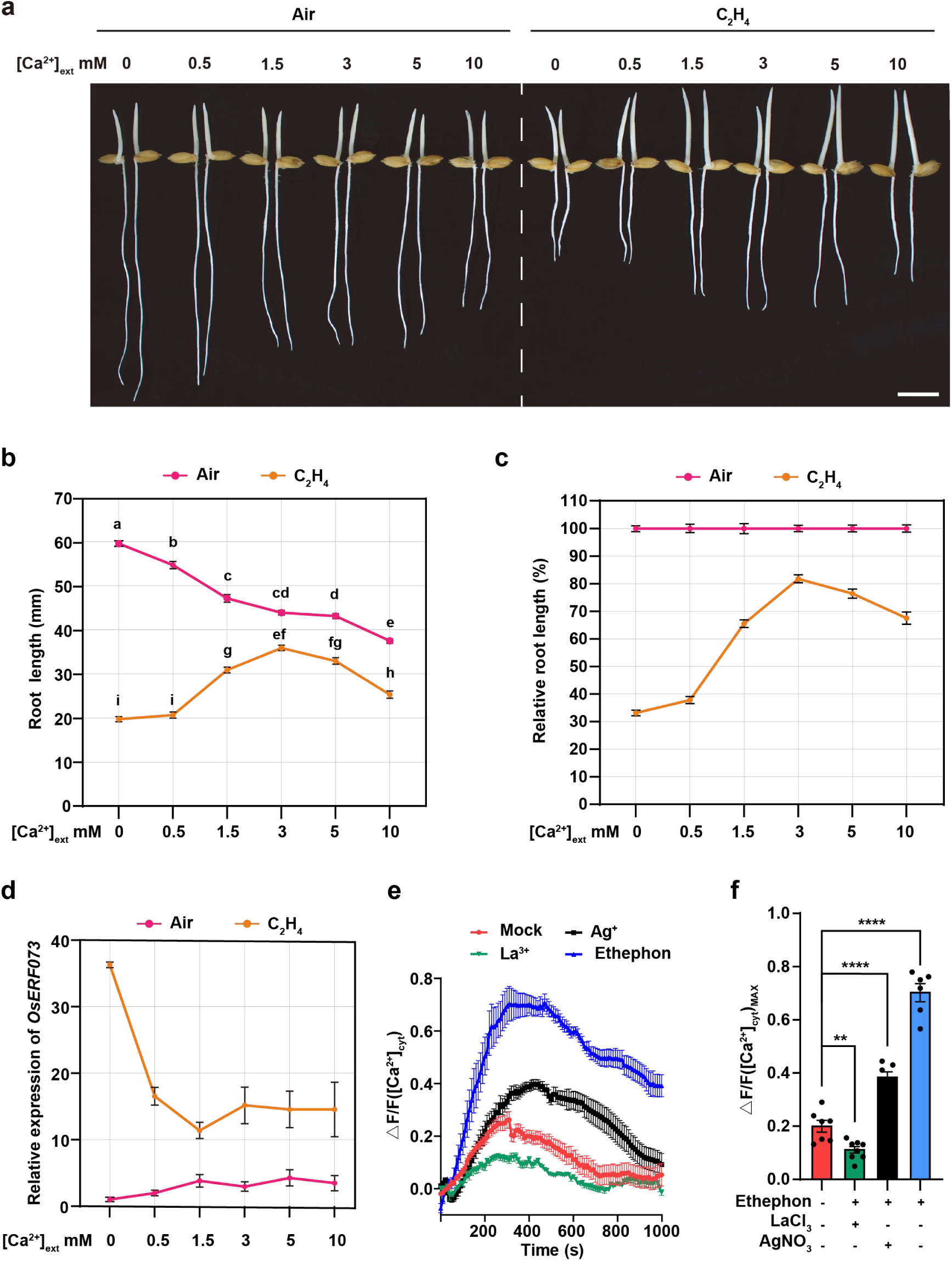
Calcium antagonizes ethylene-induced inhibition of root growth in rice. **a-c** Phenotypes of DJ rice plants treated with air of 2 ppm ethylene at different [Ca^2+^]_ext_ concentrations arranged from 0, 0.5, 1.5, 3, 5 to 10 mM, bar = 10 mm. Quantification of root length (**b**) and relative root length (**c**). Data are presented as mean ± s.e.m. and n ≥ 20 seedlings. *P* values are calculated using two-way analysis of variance (ANOVA) and multiple comparisons by the Tukey’s method. **d** Relative expression of the ethylene-responsive gene *OsERF073* in roots after 8 h treatment with/without 2 ppm ethylene. Data are mean ± s.e.m. **e, f** Ethephon-induced [Ca^2+^]_cyt_ increases in DJ callus expressing GCaMP6s (**e**) and quantification of their maximum (**f**). Data are presented as mean ± s.e.m. from three representative experiments. Before processing, the callus tissue was incubated with 0.2 mM MES (pH=4.2), 100 μM AgNO_3_, and 1 mM LaCl_3_ for 30 minutes, and then fluorescence changes were measured. 1 mM ethephon was added at 50 seconds.

Transcriptome analysis further revealed that ethylene induces the expression of WRKY transcription factors and cell wall synthesis-related genes^30^, while Ca^2+^ treatment antagonizes this induction. Many WRKY family members are associated with ethylene-mediated stress responses^31,32^, and their transcript levels were downregulated by Ca^2+^ supplementation (Supplementary Fig. 4a). Moreover, Ca^2+^ and ethylene exerted opposing effects on related biological pathways when applied individually versus in combination (Supplementary Fig. 4b). Together, these findings demonstrate a clear antagonistic relationship between calcium and ethylene signaling.

Previous studies have shown that ACC/ethephon activates calcium channels in plants, leading to an increase in the concentration of cytosolic calcium ([Ca^2+^]_cyt_)^12^. To examine whether ethylene specifically induces [Ca^2+^]_cyt_ fluctuations in rice, we monitored calcium dynamics in DJ callus expressing the calcium indicator GCaMP6s. Ethylene treatment triggered a pronounced transient increase in [Ca^2+^]_cyt_, which was abolished by La^3+^, an inhibitor of plasma membrane calcium channels (Fig. 1e, f). Meanwhile, application of Ag⁺, which disrupts Cu^+^ cofactor binding to ethylene receptors^33^, also markedly suppressed, though not completely, the ethylene-induced [Ca^2+^]_cyt_ peak (Fig. 1e, f). These findings suggest that ethylene receptors are likely involved in initiating the Ca^2+^ influx underlying these cytosolic calcium transients. We therefore focused subsequent analyses on ethylene receptors in rice.

### The ethylene receptor OsERS1 is identified as a novel calcium-permeable channel

To determine whether rice ethylene receptors have the calcium channel activity, we expressed four representative members-OsERS1 and OsERS2 (subfamily I), and OsETR2 and OsETR3 (subfamily II)^36^-in *Xenopus laevis* oocytes and performed two-electrode voltage clamp (TEVC) assays, a well-established system for characterizing plant and animal ion channels^24,25^. OsERS1 and OsERS2 elicited inward currents at-140 mV similar to the known calcium channel AtCNGC14 (cyclic nucleotide-gated channel 14) (Fig. 2a, b)^27^. However, OsETR2 and OsETR3 showed a very weak and background signal, respectively (Fig. 2a, b). Since OsERS1 localizes to the plasma membrane in rice^37^, frog oocytes, and tobacco cells (Supplementary Fig. 5), we selected it for further characterization. These data indicate that ethylene receptors in rice, especially subfamily I members, exhibit intrinsic calcium channel activity.

**Fig. 2.**
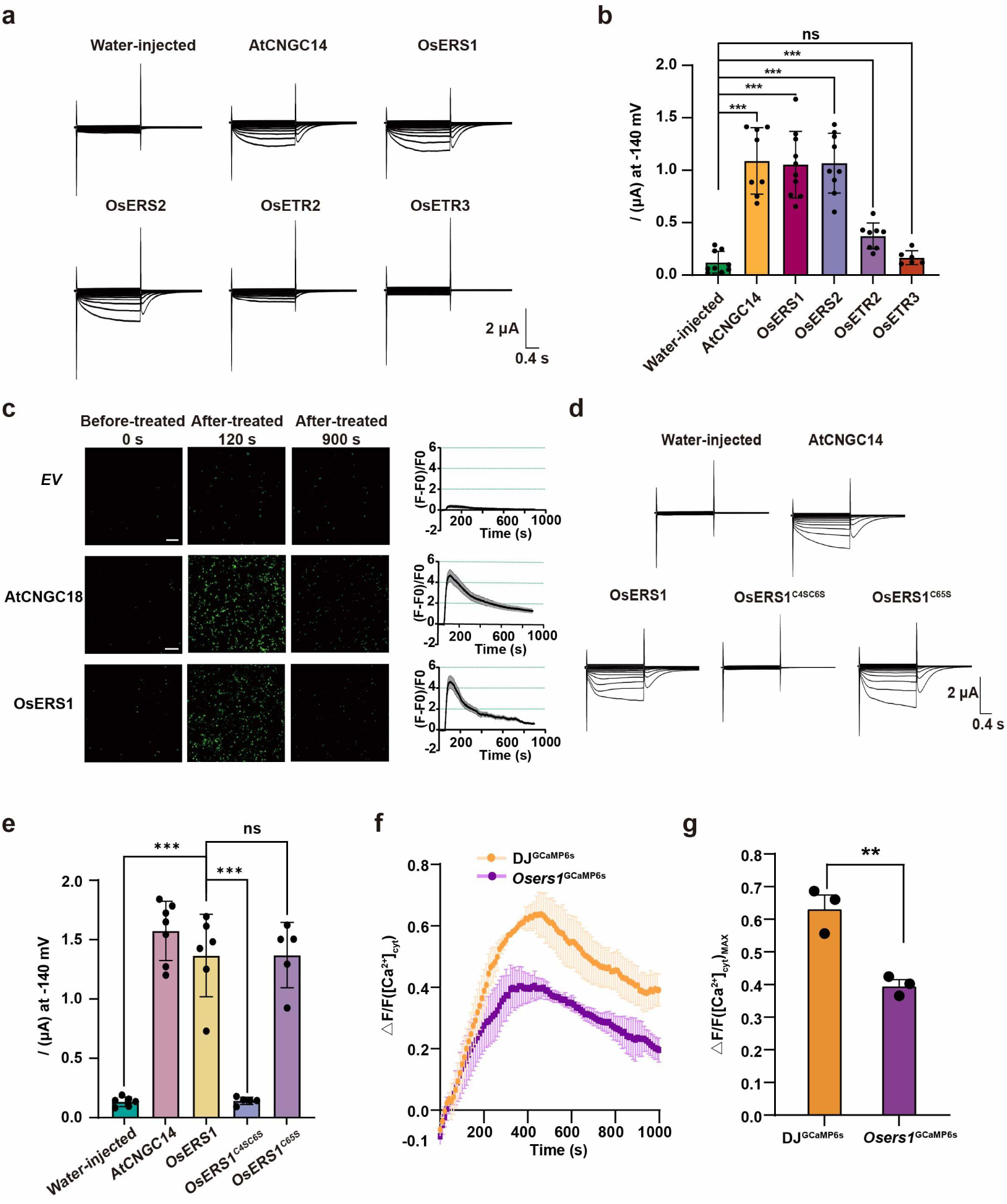
The ethylene receptor OsERS1 functions as a Ca^2+^-permeable channel. **a, b** Typical two-electrode voltage-clamp (TEVC) recordings (**a**) and current amplitudes at −140 mV (**b**) of inward currents in *Xenopus* oocytes expressing water-injected (mock), AtCNGC14 (positive control), OsERS1, OsERS2, OsETR2, or OsETR3. *P* values are calculated from two-sided Student’s *t*-tests. Data are presented as mean ± s.d., and n ≥ 6 biologically independent oocytes. **c** The plasmid expressing the target gene and the Ca^2+^ fluorescence indicator GCaMP6s was transferred into HEK293T cells, and fluorescent signals were then monitored using confocal microscopy at 5-s intervals over a 15-min acquisition period. At 60 s after initiating imaging, intracellular Ca^2+^ dynamics were triggered by adding CaCl_2_ to a final concentration of 2 mM. Data are presented as mean ± s.e.m., n > 10 independent cells. Bar = 200 μm. **d, e** Typical TEVC recordings and current amplitudes at −140 mV (**d**) of inward currents in *Xenopus* oocytes expressing water-injected (mock), AtCNGC14 (positive control), OsERS1, OsERS1^C4SC6S^, or OsERS1^C65S^. *P* values are calculated from two-sided Student’s *t*-tests. Data are presented as mean ± s.d., and n ≥ 6 biologically independent oocytes. **f, g** Ethephon-induced [Ca^2+^]_cyt_ increases in DJ and *Osers1* callus expressing GCaMP6s (**f**) and their quantification (**g**). Data are mean ±s.e.m. from three representative experiments.

We next verified the calcium permeability of OsERS1 in HEK293T cells expressing the calcium sensor GCaMP6s^38^. Upon treatment with 2 mM CaCl_2_, peak fluorescence signals from both the positive control and OsERS1-expressing cells were significantly elevated, in contrast to the undetectable signal in the negative control (Fig. 2c), indicating that OsERS1 mediates a distinct specific Ca^2+^ influx signal. Moreover, in the complementation assays of calcium channel-deficient yeast mutant^34,39^, expression of either AtCNGC14 or OsERS1 significantly rescued the growth of the *cch1mid1* mutant defective in Ca^2+^ uptake (Supplementary Fig. 6). These findings thus confirm that, in addition to its role as an ethylene receptor, OsERS1 also functions as a Ca^2+^-permeable channel.

However, this also raises a question: by what mechanism does it balance its two distinct functions as a receptor and a channel? Previous studies have identified many key sites in ETR1 of *Arabidopsis* that affect its binding to ethylene^40^. Given the high sequence conservation between AtETR1 and OsERS1 (Supplementary Fig. 7), we targeted functionally conserved critical residues: Cys4 and Cys6 (involved in protein complex formation) and Cys65 (mediating ethylene perception)^41^. When the mutation of Cys65 deficiency in ethylene binding activity, we found the calcium channel activity of OsERS1^C65S^ did not change significantly compared to OsERS1^WT^ (Fig. 2d, e), suggesting that the channel function may be independent of the receptor role. The same results were also obtained in the complementation assay of calcium channel-deficient yeast *cch1mid1* (Supplementary Fig. 6). Additionally, to test whether OsERS1 homo-complex can affect channel activity, we generated a mutant version of OsERS1^C4SC6S^ deficiency in forming disulfide bonds required for homo-dimer, tetramer, or high-molecular polymer^41^, we found that the calcium channel activity of OsERS1^C4SC6S^ was totally lost, indicating that homo-complex formation is essential for calcium channel activity of OsERS1.

The homomeric complex of OsERS1 is formed by disulfide bonds (Supplementary Fig. 8a). Luciferase complementation imaging (LCI) and co-immunoprecipitation (Co-IP) analysis demonstrated that OsERS1 undergoes homomeric protein interactions (Supplementary Fig. 8b, c). We hypothesize that homomeric complex constitute the core functional unit for both ethylene perception and channel activity. However, additional evidence is required to substantiate that protein multimerization or conformational changes underpin its functional divergence.

We observed changes in ethylene-induced elevation of calcium peaks in mutants. The ethylene-sensitive Ca^2+^ peak was significantly reduced by 37% in *Osers1* (Fig. 2f, g). These data indicate that ethylene-induced [Ca^2+^]_cyt_ elevation is at least partially mediated by the calcium channel OsERS1. The residual [Ca^2+^]_cyt_ fluctuations observed in *Osers1* mutants likely result from functional redundancy among ethylene receptors.

In conclusion, the channel activity of OsERS1 is dependent on specific residues critical for homomeric complex formation, and this functional property is distinct from its canonical role in ethylene perception. OsERS1 possesses dual functions as an ethylene receptor and a calcium channel, with regulatory roles in rice growth and development.

### The OsERS1 channel is permeable to both mono- and di-valent cations

We further systematically analyzed the channel characteristics of OsERS1. When oocytes expressing OsERS1 were perfused with increasing external Ca^2+^ concentrations, the currents mediated by the OsERS1 channel gradually increased, and this trend exhibited a strong positive correlation (Fig. 3a). These results indicate that OsERS1 channel activity is regulated by and dependent on external Ca^2+^ concentrations. Additionally, La^3+^ significantly suppressed the OsERS1-mediated current to background levels, comparable to the negative control (Fig. 3b), demonstrating that La^3+^ completely inhibits OsERS1 channel activity.

**Fig. 3.**
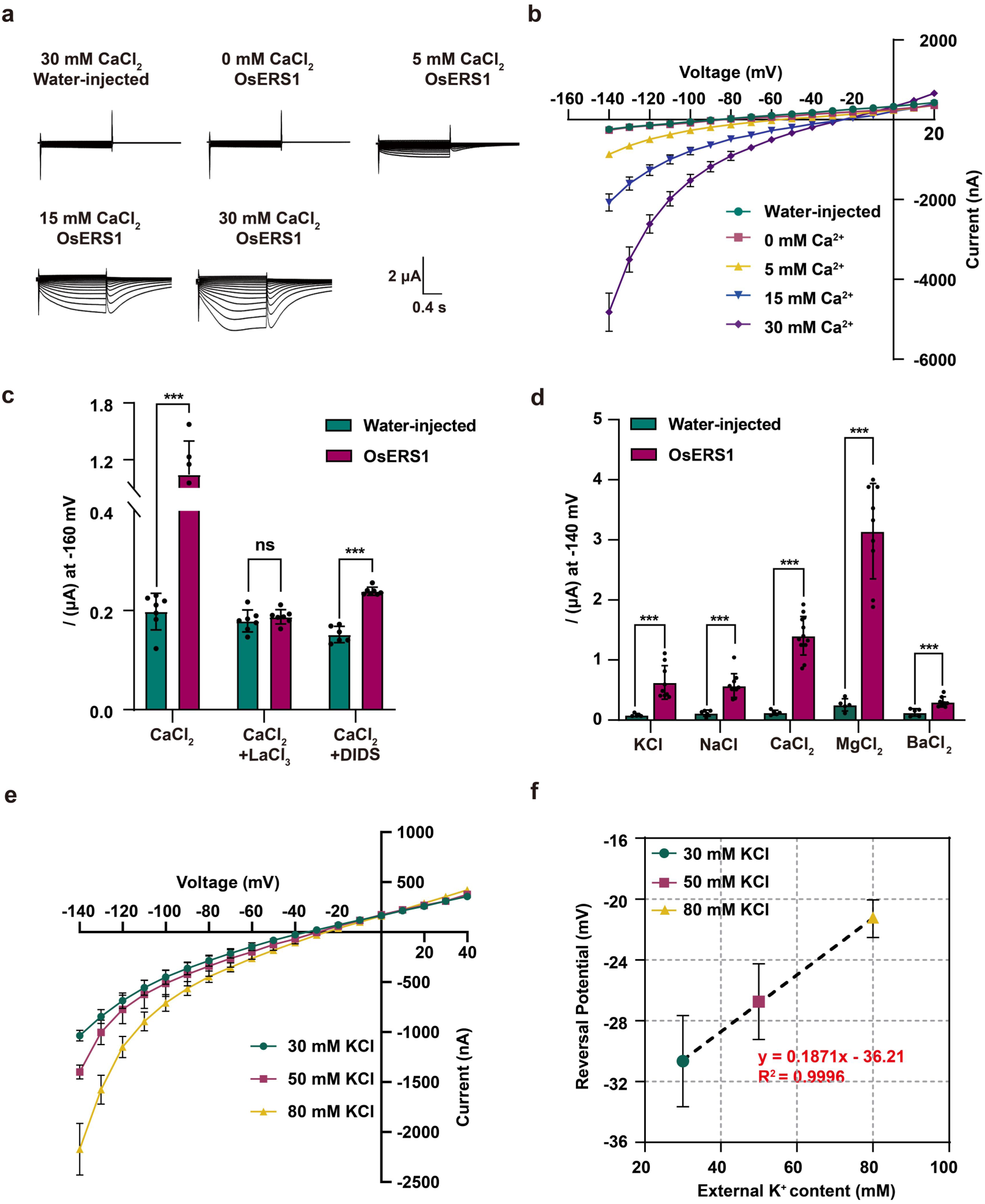
OsERS1 is a canonical cation channel. **a, b** Typical TEVC recordings (**a**) and current-voltage relationships (**b**) of inward currents in oocytes expressing OsERS1 perfused with indicated extracellular Ca^2+^ concentrations. Data are presented as mean ± s.e.m., and n ≥ 6 biologically independent oocytes. **c** The currents at -160 mV were recorded in oocytes expressing OsERS1 perfused with 30 mM CaCl_2_, 30 mM CaCl_2_ + 1 mM LaCl_3_, or 30 mM CaCl_2_ + 1 mM DIDS by TEVC assay. Data are presented as mean ± s.d., and n ≥ 6 biologically independent oocytes. *P* values are calculated from two-sided Student’s *t*-tests. **d** The currents at -140 mV were recorded in oocytes expressing OsERS1 perfused with 30 mM KCl, 30 mM NaCl, 15 mM CaCl_2_, 15 mM MgCl_2_ or 15 mM BaCl_2_ by TEVC assay. Data are presented as mean ± s.d., and n ≥ 5 biologically independent oocytes. *P* values are calculated from two-sided Student’s *t*-tests. **e. f** Current-voltage relationships of inward currents (**e**) and reversal potentials (**f**) in oocytes expressing OsERS1 perfused with varying extracellular KCl concentrations. Dashed line is the least-square fit (**f**). Data are presented as mean ± s.e.m., and n = 3 independent oocytes.

We treated *Xenopus* oocytes with DIDS (4, 4’-Diisothiocyanatostilbene-2, 2’-disulfonate), which blocks Cl^-^ channels known to amplify Ca^2+^ signals in TEVC^42,43^, resulted in a significantly enhanced primary Ca^2+^ signal relative to negative controls (Fig. 3c). This finding further supports that OsERS1 functions as a Ca^2+^-permeable channel. To further characterize the ion selectivity of OsERS1, we recorded current signals in perfusion solutions supplemented with distinct ions (Fig. 3d). The results demonstrated that the OsERS1 can transport not only monovalent ions like K^+^ and Na^+^, but also divalent ions like Ca^2+^, Ba^2+^and Mg^2+^. This demonstrates that OsERS1 channel can permeate cations like animal TRP (transient receptor potential) channels^35^.

Our experimental results show that OsERS1 is a non-selective cation channel. To further validate this conclusion, we measured the reversal potential (*E*rev) of the cation channel via TEVC assay. By examining the impact of K⁺ concentration ([K^+^]), we got a linear relationship between the *E*rev and the external [K^+^], and reduction of extracellular [K^+^] from 80 mM to 30 mM shifted *E*rev by 10 mV, suggesting that the *E*rev of OsERS1 channel depends on [K^+^] (Fig. 3e, f), which is sufficient to prove that the ethylene receptor is an ion channel.

All the above experimental results confirm that OsERS1 exhibits the characteristics of an ion channel, indicating that OsERS1 itself is an ion channel. However, the relationship between its receptor function and channel function requires further investigation. Taken together, these results demonstrate that the ethylene receptor OsERS1 is permeable to both mono- and di-valent cations. This finding reinforces its typical channel function, which is functionally distinct from its classical receptor role, uncovering a previously unrecognized dual identity.

### The physiological interplay of calcium and ethylene signals requires OsERS1 and OsERS2

We analyzed the phenotype of the mutants and the culture conditions were referenced to previous treatments^29^. The phenotype of ethylene-inhibited root elongation was restored with the increase of extra Ca^2+^ concentration from 0 to 1.5 and 3 mM, consistent with the aforementioned CAER effect. However, in *Osers1* mutants, there was almost no CAER effect on the phenotype change of rice roots treated with ethylene and gradient Ca^2+^ (Fig. 4a, b). These data indicate that the channel activity of OsERS1 plays a significant role in inhibiting the role of the classical ethylene signaling pathway, thereby genetically establishing a direct interplay between Ca^2+^ and ethylene signaling in rice. As a member of the same family, *Osers2* mutants exhibit a similar CAER effect to *Osers1* mutants (Fig. 4c, d). These observations imply that the dual receptor-channel function of ERS1 and ERS2 can effectively coordinate the close crosstalk between calcium and ethylene signals. The receptor function and channel function may be in an antagonistic relationship, and such antagonism becomes more pronounced under ethylene stress. This suggests that under ethylene stress, plants can counteract the stress by regulating the functions of ERSs.

**Fig. 4.**
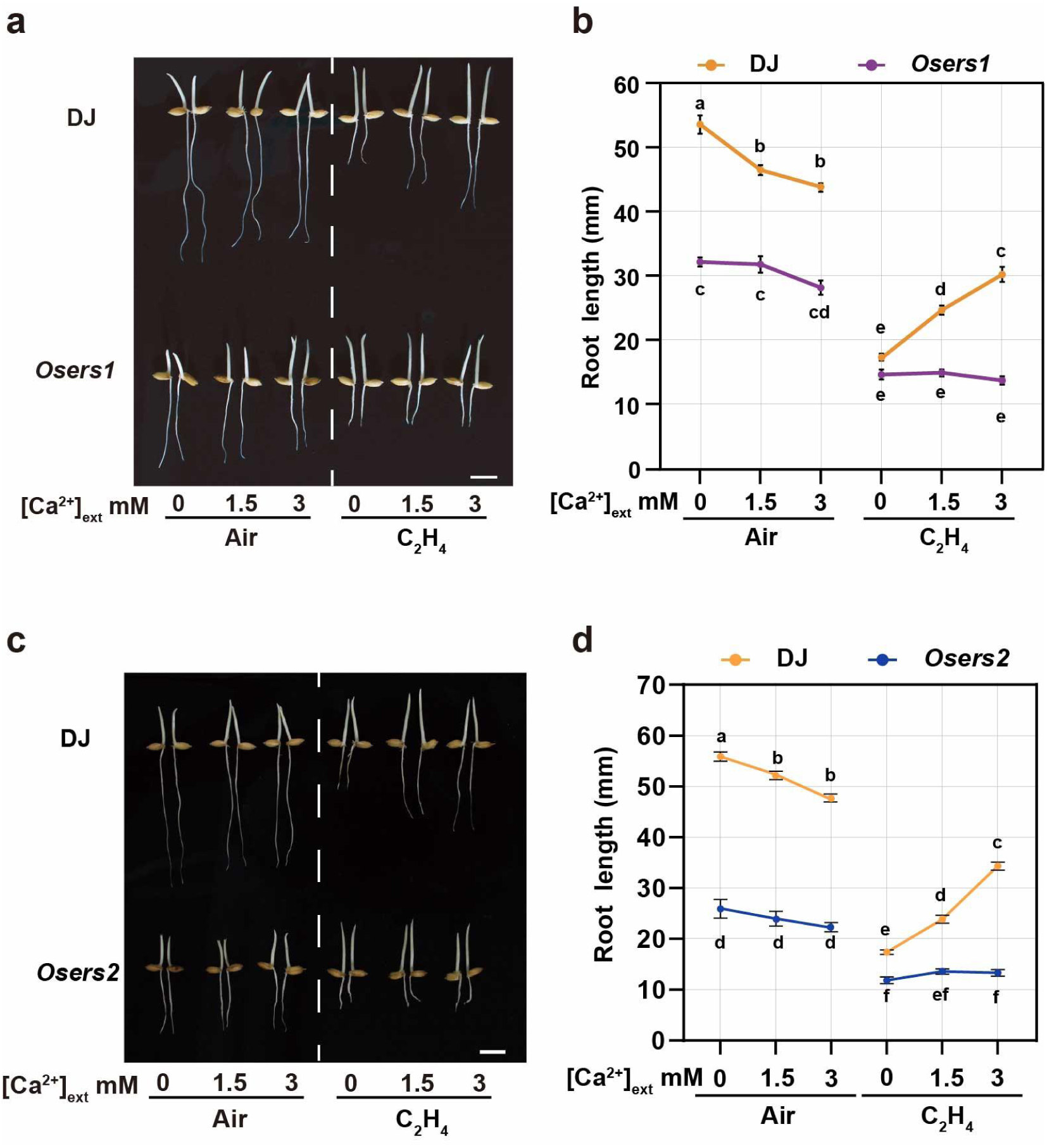
The physiological interplay between calcium and ethylene signals is dependent on OsERS1 and OsERS2. **a, b** Phenotypes of *Osers1* treated with air of ethylene at different extra Ca^2+^ ([Ca^2+^]_ext_) concentrations arranged from 0, 1.5, and 3 mM, bar = 10 mm. Quantification of root length (**b**). **c, d** Genetic phenotype of the *Osers2* mutant. bar = 10 mm. Quantification of root length (**d**). Data are mean ± s.e.m., and n ≥ 20 seedlings. *P* values are calculated using two-way analysis of variance (ANOVA) and multiple comparisons by the Tukey’s method.

Based on our experimental findings, we propose that under ethylene stress, when calcium concentration is low, ethylene binds to OsERSs, activating the ethylene signaling pathway and inhibiting rice root growth; In contrast, under high-calcium conditions, calcium antagonizes ethylene action. OsERSs exert their channel function, the ethylene signaling pathway is perturbed, and rice root length is partially restored. Consequently, this study marks the identification of a hormone receptor-type ion channel, which has not been reported in the hormone receptors of all living organisms.

## DISCUSSION

Here, we show that OsERS1 functions as Ca^2+^-permeable cation channel, and thereby exert a regulatory effect on rice root growth. Notably, the channel and receptor activities exist as independent functional modules within the same OsERS1 protein molecule. Consequently, this study uncovered a hormone receptor-type channel, marking a paradigm-shifting discovery in hormone signal transduction. As well, this finding could serve to optimize crop root system architecture, and improv the efficiency of water and fertilizer utilization—thereby increasing crop yield or facilitating adaptation to diverse planting environments.

In animals, certain receptors such as TRP channels function as both receptors and ion channels. For instance, TRPV1 receptors can be activated by capsaicin and act as calcium/ion channels^52^. Upon neurotransmitter binding, these receptors induce conformational changes in channel proteins, leading to ion channel opening or closure. This alters plasma membrane ion permeability, converts extracellular chemical signals into electrical signals, and subsequently modulates postsynaptic cell excitability to elicit biological responses^47,48^. However, no hormone receptor has been reported to function as an ion channel across organisms to date. Here, we present the first example of a hormone receptor-type ion channel (HRIC), establishing this new concept, expanding the receptor-ion channel family, and broadening its scope.

In ethylene-mediated physiological responses, where OsERS1 functions dually as an ethylene receptor and a calcium channel; however, the mechanistic intricacies underlying their interplay remain to be deciphered. The TRP family members TRPM6 and TRPM7 are bifunctional transmembrane proteins that possess both ion channel and kinase domains^53,54^. Ethylene receptors share evident sequence homology with bacterial and eukaryotic two-component signaling kinases. It has been reported that rice ethylene receptor proteins possess a diverged histidine kinase domain and have been proven to exhibit serine/threonine kinase activity, which can phosphorylate their own receiver domain^55^. Future studies should focus on exploring the intricate regulatory interactions between the receptor, channel and kinase activities of ethylene receptors.

Ca^2+^ signaling serves as a central hub regulating diverse physiological processes in plants. Activation of calcium channel of the ethylene receptors tightly couples ethylene signals to these processes, enabling precise spatiotemporal control over growth and developmental programs^13,14,17,38^.

Although we have identified that OsERS1 performs a channel function in addition to its role as a receptor, and have experimentally validated this function, the specific mechanism underlying the integration and differentiation of these two functions need further investigation. Additionally, the role of OsERS1’s channel activity in the classical ethylene signaling pathway remains to be elucidated. These unresolved questions not only highlight key directions for future studies to dissect the dual-function regulation of OsERS1 but also hold the potential to refine and expand the “hormone receptor-type ion channel” paradigm.

## Methods

### Plant Materials and Growth Conditions

The *OsERS1* (LOC_Os03g49500) T-DNA knockout mutant *Osers1* (PFG_1B-08531.L), *Osers2* (LOC_Os05g06320) are in the Dongjin (DJ) background. The homogenous *Osers1* and *Osers2* were identified by PCR (The primer sequences are in Supplemental Table 1). The ethylene treatments were performed as previously described^29^. Seeds were soaked in tap water and germinated at 37°C in the dark for 2 days. Then cultured at 28°C in the dark.

### Root elongation assay

Place the germinated seeds on a wire mash support and place them in an airtight container. Ethylene gas (2 ppm) was injected into the boxes using a syringe. The seedlings were cultured at 28°C in the dark for 3 days with different concentrations of CaCl_2_. Images were captured using a Canon EOS 550D camera, Root elongation was quantified by measuring the linear distance between the initial root tip mark and the terminal tip using ImageJ software. The statistical method for relative root length is defined as follows: the root length of wild-type rice without additional calcium supplementation is set as 100%, and then compared with the root lengths of rice under other conditions or of mutant rice.

### Gene Expression Analysis Using RT-PCR

Three-day-old etiolated seedlings were treated for up to 8 h with 2 ppm ethylene or air or with different concentrations of CaCl_2_ with or without ethylene. After treatment, the shoots and roots were harvested and immediately frozen in liquid nitrogen. Total RNA was extracted by Trizol (LABLEAD, R1000), and 1 μg total RNA was used for reverse transcription (LABLEAD, F0202). The obtained cDNA was used as a template to detect the expression of the interest genes by qPCR assay (LABLEAD, R0202-02). *OsActin2* was used as the internal control and relative expression of target genes was calculated using the 2(*-ΔΔCt)* method. The primers sequences are listed in Supplemental Table 1.

### Two-electrode voltage clamp recording from *Xenopus* oocytes

The full-length coding sequences (CDS) of *AtCNGC14, OsERS1, OsERS2, OsETR2*, and *OsETR3* were cloned into the expression vector *pGEMHE* via homologous recombination and expressed in *Xenopus* oocytes. Target genes were amplified from the recombinant plasmids, and capped RNA (cRNA) was synthesized using the mMESSAGE mMACHINETM T7 Transcription Kit (Thermo Fisher Scientific, AM1344) following the manufacturer’s protocol.

The oocytes were isolated and incubated overnight at 16°C in calcium-free ND96 solution (96 mM NaCl (Sigma-Aldrich, S3014), 2 mM KCl (Sigma-Aldrich, P9333), 1 mM MgCl_2_ (Sigma-Aldrich, G3068), 10 mM HEPES ((Sigma-Aldrich, H3375)/ NaOH (Sigma-Aldrich, S5881), pH 7.4) supplemented with 50 μg/mL gentamicin. Each oocyte was microinjected with 23 nL of cRNA (1000 ng/μL) and then transferred to complete ND96 medium containing 1.8 mM CaCl_2_ for further culture at 16°C for 48 h.

Oocytes were voltage-clamped using a TEV-200A amplifier (Dagan) and monitored by computer using a Digidata 1440 A converter and pCLAMP 10.4 software (Axon Instruments). Electrodes (filled with 3 M KCl) had resistances of 0.5-1.5 MΩ, and the membrane potential was clamped at -30 mV. The perfusion solution for calcium channel activity assays contained 30 mM CaCl_2_, 2 mM KCl, 1 mM MgCl_2_, and 10 mM MES (Sigma-Aldrich, M3671)/Tris (pH 5.6), with osmolarity adjusted to 220-230 mOsm/kg using D-mannitol (Sigma-Aldrich, M1902). Ion selectivity experiments utilized MES/Tris buffer (pH 5.6) containing 15 mM divalent cations (Ca^2+^, Ba^2+^, Mg^2+^) or 30 mM monovalent cations (Na⁺, K⁺). Pharmacological tests included 1 mM DIDS (anion channel inhibitor) or 1 mM LaCl_3_ (calcium channel blocker). Voltage-step protocols were applied from +20 mV to -140 mV in -10 mV increments, with each pulse lasting 2 seconds. Data were analyzed using Origin 2019b and Clampfit 10.7.

### Calcium Imaging in HEK293T Cells

The full-length coding sequences of *AtCNGC14, OsERS1, OsERS2, OsETR2,* and *OsETR3*, were directionally cloned into the pBudCE4.1-GCaMP6s dual-expression vector downstream of the CMV promoter using *Hin* d III/*Xba* I restriction sites to construct recombinant plasmids co-expressing calcium indicators^38^. HEK293T cells were cultured in high-glucose DMEM medium (CellMax, CGM102.05) supplemented with 10% fetal bovine serum (CellMax, SA211.02) and 1% penicillin-streptomycin (Solarbio, P1400) at 37°C under 5% CO_2_. Cells were inoculated into dishes containing coverslips (NEST, 801007) 24 h before transfection. Plasmid DNA was purified using the Endotoxin-Free Plasmid Mini Kit (TIANGEN, DP118-02) and mixed with PEI 25K (Polysciences, 23966-100) in high-glucose DMEM, incubated for 15 minutes, and transfected into cells. Complete medium replacement was performed 6 h post-transfection, followed by 48 h incubation. Confocal imaging (Leica SP8) utilized 488 nm excitation and 505-521 nm emission parameters, with images captured every 5 s for 180 frames using Leica LAS X software. Cells were perfused with a buffer (pH 7.4) containing 120 mM NaCl, 3 mM KCl, 1 mM MgCl_2_, 1.2 mM NaHCO_3_, 10 mM glucose, 10 mM HEPES, and 2.5 mM EGTA (Sigma-Aldrich, 03777). After 60 seconds of baseline recording, 2 μM CaCl_2_ was added to activate channel-mediated calcium influx. The fluorescence change ratio *(ΔF/F)* was calculated as *ΔF/F = (F − F₀)/F₀*, where *F₀* represents baseline fluorescence at rest. Time-dependent fluorescence curves were analyzed using GraphPad Prism 10.

### Complementation assay of calcium channel-deficient yeast

Full-length coding sequences of *AtCNGC14*, and *OsERS1* were cloned into the yeast expression vector YEP124 by homologous recombination. Functional complementation assays were performed using the BY4741-derived calcium channel-deficient yeast mutant strain (*cch1 mid1*). Yeast cells were grown to logarithmic phase (OD_600_ = 0.8) in YPD liquid medium (1% Yeast Extract (OXOIP, LP0021B), 2% Peptone (OXOIP, LP0042B), 2% Glucose) containing 2% glucose under 30°C with shaking. A 1.5 mL aliquot of culture was centrifuged at 3000×g for 5 min, and 100 μL of 0.1 mol/L LiAC (Sigma, 517992) was added to each tube for re-suspension, and then incubated at 30°C for 10 min to prepare yeast component cells. The plasmid containing the interest gene was transformed into yeast cch1 mid1 by 50% PEG4000 (Sigma-Aldrich, 81240). After recovery at 28°C for 3 h, yeast cells were collected and plated on SD/-His-Leu-Ura (Coolaber, PM2170) solid medium for selection. The correct colony was selected and the wild-type BY4741 colony was cultured with YPD medium to OD600 = 3.0. 100 μL aliquot of yeast extract was mixed with 4 ml of sterilized top agar medium (20 g/L yeast extract, 20 g/L tryptone, 20 g/L glucose, 0.7% agar) and overlaid onto YPD solid medium. Place around filter paper on the medium, then add 10 μL of 10 mg/ml α-factor (Macklin) to the center of the filter paper. Incubate at 28°C for 12 h to observe the growth of yeast. Regarding quantification, measure the grayscale value of the circle formed around the background and filter paper, with the background grayscale value as 1, and quantify the relative grayscale value of the circle.

### Calcium Imaging in rice callus

The husked rice seeds were sterilized by immersion in 70% ethanol for ∼2 min, followed by 15% sodium hypochlorite solution for 30 min with shaking, and rinsed three or four times with sterile water on an ultraclean workbench. N6 was used for callus induction and subculture. The pH of the medium was adjusted to 5.8 with 5 M KOH, and 3 g/L of Phytagel was added before autoclaving at 121°C for 20 min. Seeds were incubated in induction medium for 7 days at 28°C in the light. Vector transformation and material screening refer to article^55,56^. Successfully transformed callus was picked out under a fluorescence microscope 24 h in advance, gently dispersed, and placed on N6 medium, and the next day, they were left to stand for 10-15 min under 488 nm light (blue light activates the calcium signal) before recording changes in the fluorescence signal. For the addition of inhibitors (0.2 mM MES (pH=4.2), 100 μM AgNO_3_, and 1 mM LaCl_3_), incubate for 30 min before testing. 1 mM ethephon treatment is added at 50 seconds. Image J (v.1.51j8) was used to analyze GCaMP6s signals over time at several regions of interest in the callus. To calculate the fractional fluorescence change (*△F/F*), the equation *△F/F=(F-F0)/F0* was used, where F denotes the average baseline fluorescence determined by the average of F over the first 50 frames of the recording before the treatment.

### Co-immunoprecipitation (Co-IP) assay

Co-IP assays in this study were based on^57^. For coimmunoprecipitation of OsERS1, constructs containing OsERS1-FLAG and OsERS1-GFP were co-transformed into *N. benthamiana* leaves. The protoplasts were incubated at 22 ℃ for 16 hours in the dark, and then the transformed leaves were collected for the Co-IP assay. Total protein was extracted with IP buffer [150 mM Tris-HCl (pH 7.5), 50 mM NaCl, 10% glycerin, 2% (v/v) protease inhibitor cocktail (Sigma)]. Incubate the samples on ice for 20 min, vortexing every 5 min. The 10% supernatant was obtained by centrifuging twice at 16,000×g for 10 min at 4°C, 100000×g at 4°C for 1 h. 1% DDM was used in the IP buffer to lyse cell membranes for about 2-3h. Subsequently, the beads (Anti-DYKDDDDK (Flag) Affinity Gel) were washed 5 times at 4°C with washing buffer [150 mM Tris-HCl (pH 7.5), 50 mM NaCl, 10% glycerin, 0.03% DDM, 1% (v/v) protease inhibitor cocktail]. Then, it was added to the lysed supernatant and incubated for 3 h at 4°C with rotations. Elution proteins are competed with 3 x FLAG peptide (F1001, LABLEAD). The eluted immunoprecipitates were immunoblotted with anti-GFP, anti-FLAG.

### Luciferase complementation imaging (LCI) assay

The LCI assay is used to study protein-protein interactions. The *Cluc-OsERS1*, *Cluc-OsERS1^C4SC6S^*, *Cluc-AtRBOHD*, *OsERS1-Nluc*, *Nluc-OsERS1^C4SC6S^*, *AtBIK1-Nluc*, and *AtBAK1-Nluc* constructs were transformed into *N. benthamiana* leaf cells through A. *tumefaciens* strain GV3101. Transformed *N. benthamiana* plants were grown in a greenhouse at 22℃ under a 15-h light/9-h dark photoperiod. LCI assay images were captured using a low-light, cooled charge-coupled device imaging apparatus.

### Subcellular localization analysis

To observe the subcellular localization of OsERS1, *N. benthamiana* leaves were used for expression of GFP-OsERS1 proteins, and fluorescence was detected using confocal laser scanning microscopy (Leica SP8). The coding region of OsERS1 was trans located to the *1300-35S::EGFP* vectors. Purified plasmids were then transformed into tobacco leaves as described in the Co-IP assay. After incubation at 25°C under a 16-h-light/8-dark-h photoperiod for 2 d, fluorescence signals were detected using Leica SP8.

For the subcellular localization of OsERS1 in *Xenopus* oocytes, the cRNA of OsERS1 synthesized via *in vitro* transcription using the pGEMHE-EGFP expression vector, was microinjected into oocytes. After 48 h of expression, protein levels were analyzed by Leica SP8. GFP fluorescence was excited at 488 nm, and at least 20 randomly selected membrane-localized fluorescent regions per oocyte were quantified using Leica LAS X software to calculate the mean fluorescence intensity as a measure of protein expression. Uninjected oocytes served as negative controls to account for endogenous fluorescence interference.

### Transcriptome analysis

To analyze the ethylene-responsive genes in roots, total RNA was isolated from the root tips of rice plants using TRIzol reagent (Ambion, 15596018) according to the manufacturer’s instructions. Two biological replicates were performed. Raw reads were filtered with fastp, and aligned to the reference genome using hisat2^58^. The resulting sam file containing mapped reads were converted to the bam format by SAM tools. After that, gene counts were called from the bam files using feature Counts. The R package DESeq2 was used to retrieve differentially expressed genes^59^. Results were filtered with a threshold |log_2_(fold change)|>1, P value<0.05. After that, Gene Ontology analysis was performed using the R package cluster-Profiler.

## Acknowledgments

We are grateful to Jin-Song Zhang from the Institute of Genetics and Developmental Biology, Chinese Academy of Sciences for kindly sharing the *ers1* and *ers2* seeds. This work was supported by the Program of Natural Science Foundation of China (No. 32222011 and 32170580 to L.Y.L), the Capacity Building for Sci-Tech Innovation-Fundamental Scientific Research Funds (No. 19530050165 to L.G.L.), Beijing Natural Science Foundation (5222001 to L.Y.L)

## Author contributions

L. L. and L.-G. L., conceived the project and designed the research. Z. Y., Z. Y., C. L., Y. C., E. Y., C. S., Z. Y., S. L., and H. T. performed the experiments, Z. Y., Z. Y., C. L., Y. C., C. S., Z. Y., S. L., E. Y., H. T., D.K., L. L., and L.-G. L. participated in the data analysis. L. L., and L.-G. L. wrote the manuscript with contributions from all authors. All authors contributed to the article discussed the results and have approved the submitted version.

## Supplemental figures

**Supplementary Figure 1.**
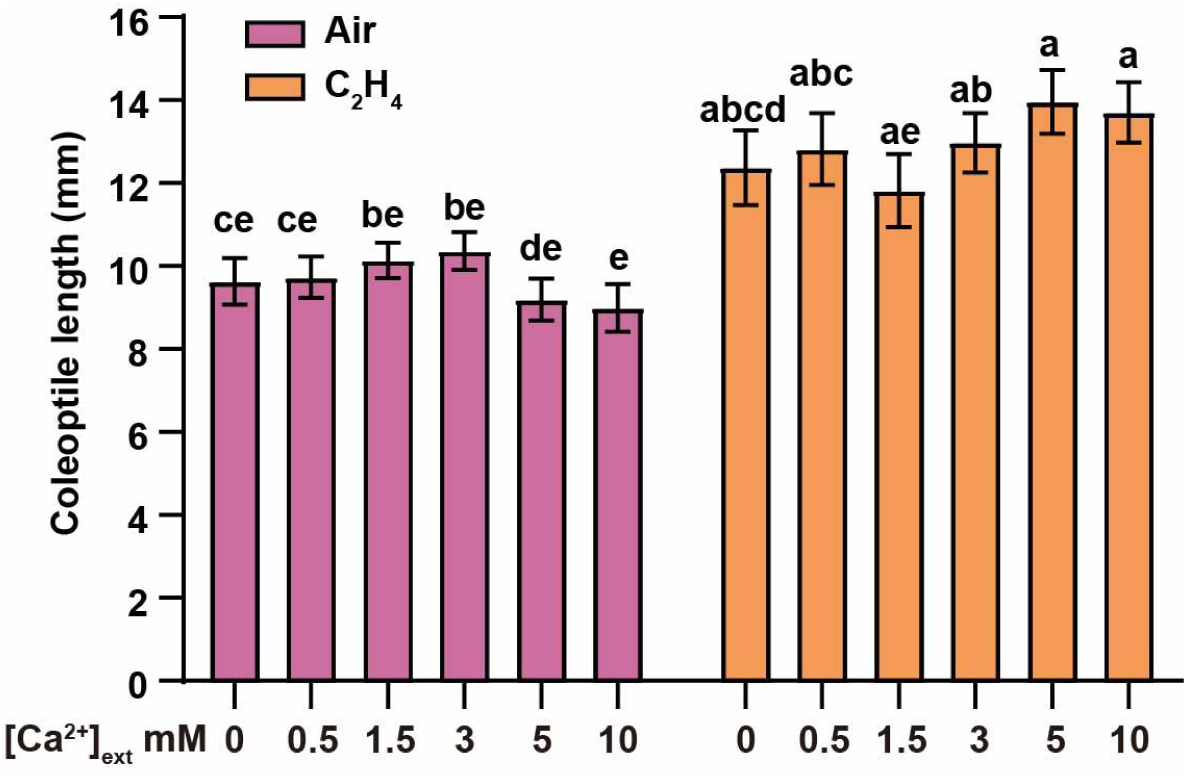
The effect of calcium on coleoptile growth under air or ethylene in rice. Coleoptile lengths of rice seedlings exposed to varying calcium concentrations in air or C_2_H_4_ atmospheres. Data are presented as mean ± s.e.m. and n ≥ 20 seedlings. *P* values are calculated using two-way analysis of variance (ANOVA) and multiple comparisons by the Tukey’s method.

**Supplementary Figure 2.**
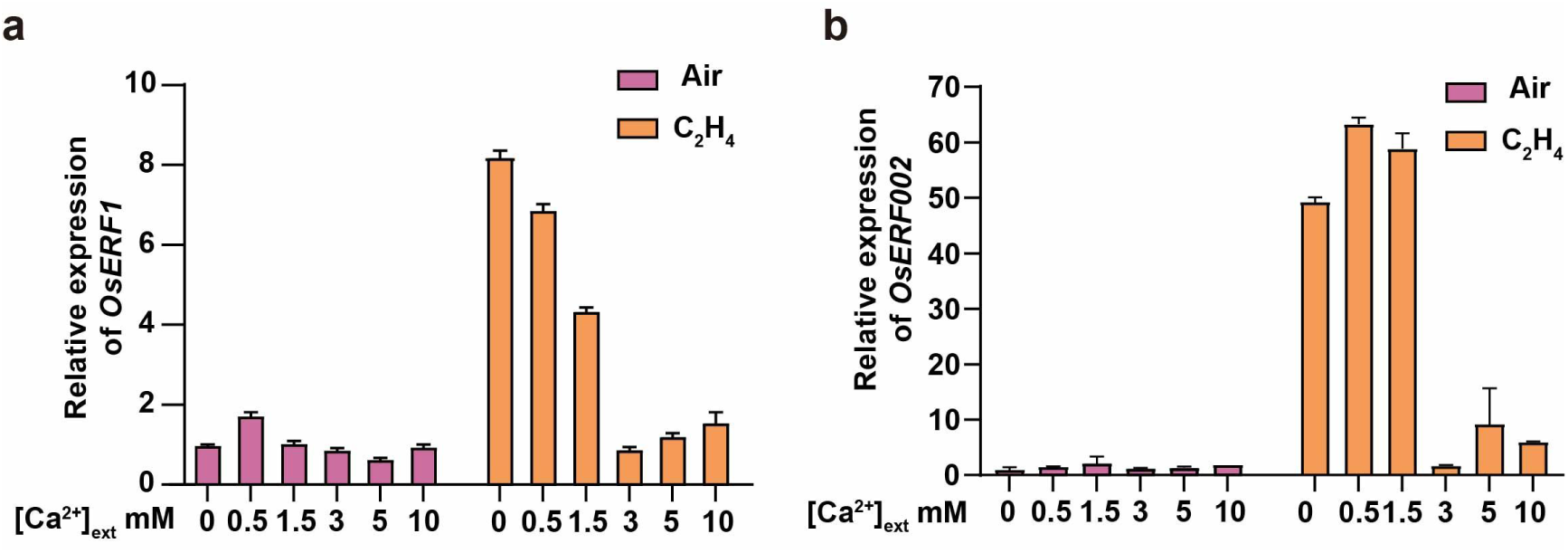
Relative expression levels of ethylene-responsive genes. **a, b** Calcium-modulated ethylene-responsive gene expression of *OsERF001*(**a**) and *OsERF002* (**b**) in WT. Three-day-old dark-grown seedlings were treated with or without 2 ppm ethylene for 8 h, and the RNA was extracted for qRT-PCR. Actin2 was used as the loading control.

**Supplementary Figure 3.**
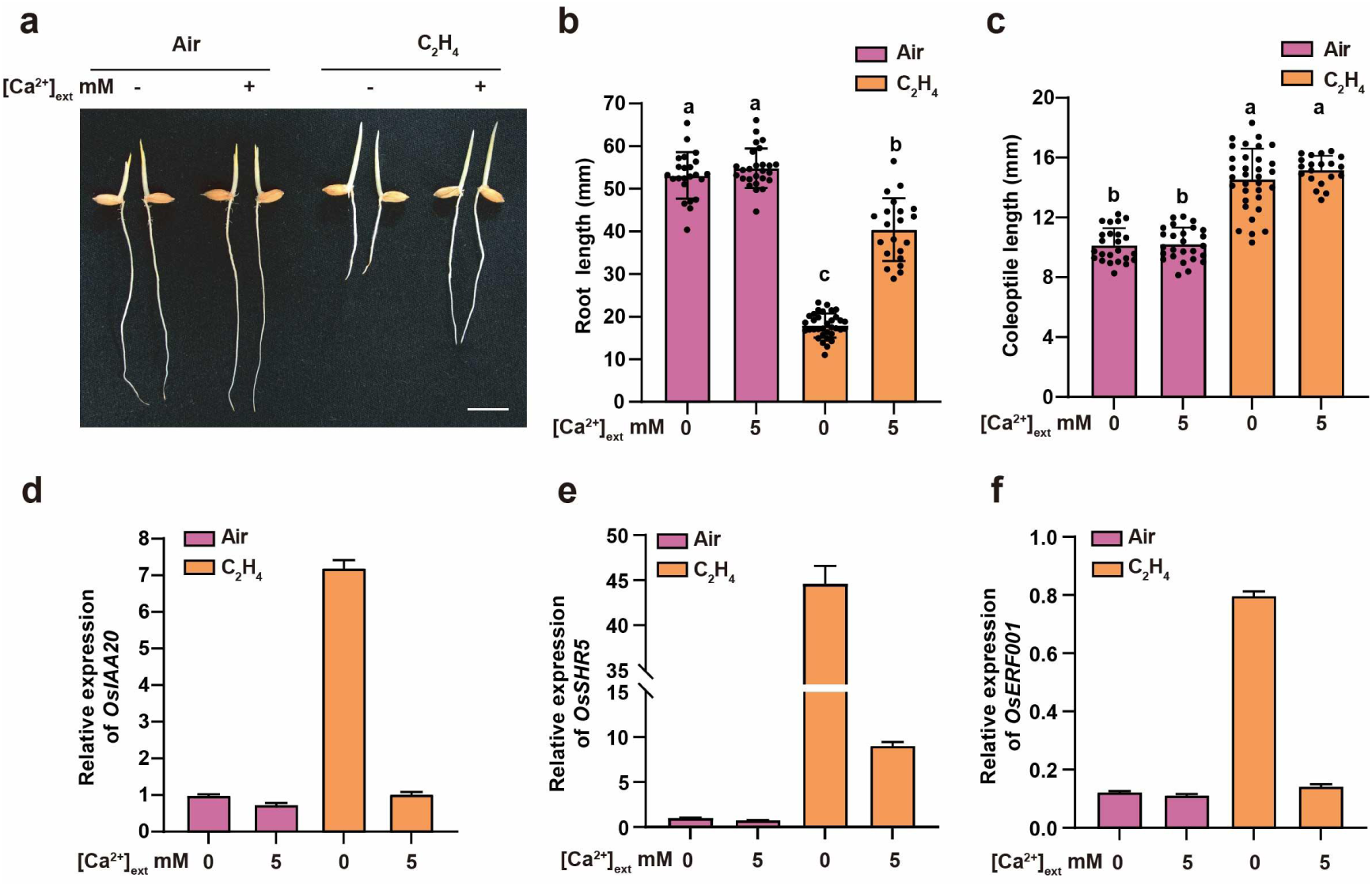
The effect of Ca^2+^ on root growth and ethylene response in rice Nipponbare. a, Phenotypes of DJ rice plants in dark with or without containing 5 mM [Ca^2+^]_ext_ (**a**) in air or 2 ppm ethylene, Bars = 10 mm. **b,** Quantification of root length. **(c)** Quantification of coleoptile length. **d-f,** Relative expression level of ethylene-responsive genes that were preferentially induced by ethylene in the roots. Three-day-old dark-grown seedlings were treated with or without 2 ppm ethylene for 8 h, and the RNA was extracted for qRT-PCR. Actin2 was used as the loading control.

**Supplementary Figure 4.**
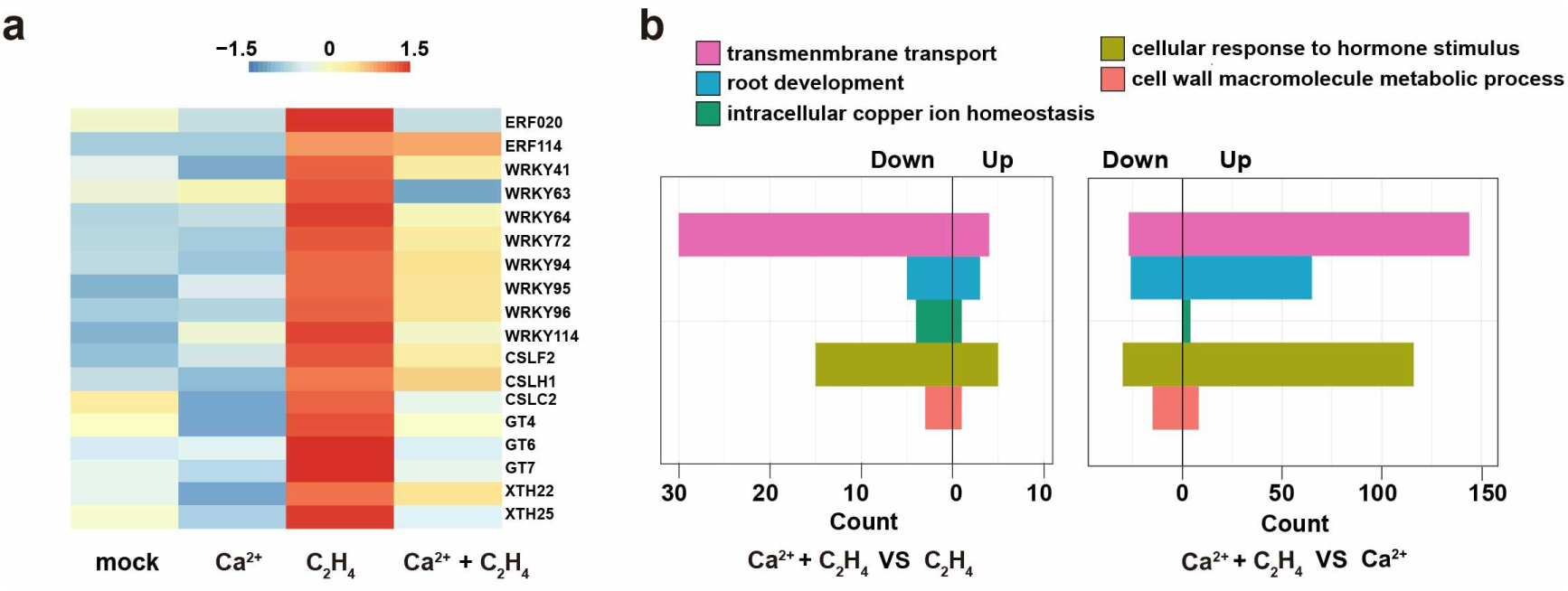
Gene Ontology enrichment analysis of ethylene-responsive genes in roots. **a, b** Gene Ontology enrichment of ethylene-responsive genes in root analyzed by RNA-seq. Heatmaps showing expression z-scores of WRKY family and cell wall synthesis-related genes (**a**). GO functional enrichment of differentially expressed genes (DEGs) (**b**). The downregulated genes are shown in left (log_2_(fold change) < -1, P < 0.05), and the upregulated genes are shown in right (log_2_(fold change) > 1, P < 0.05). Two biological replicates were analyzed. DEG, differentially expressed gene; PCA, principal component analysis.

**Supplementary Figure 5.**
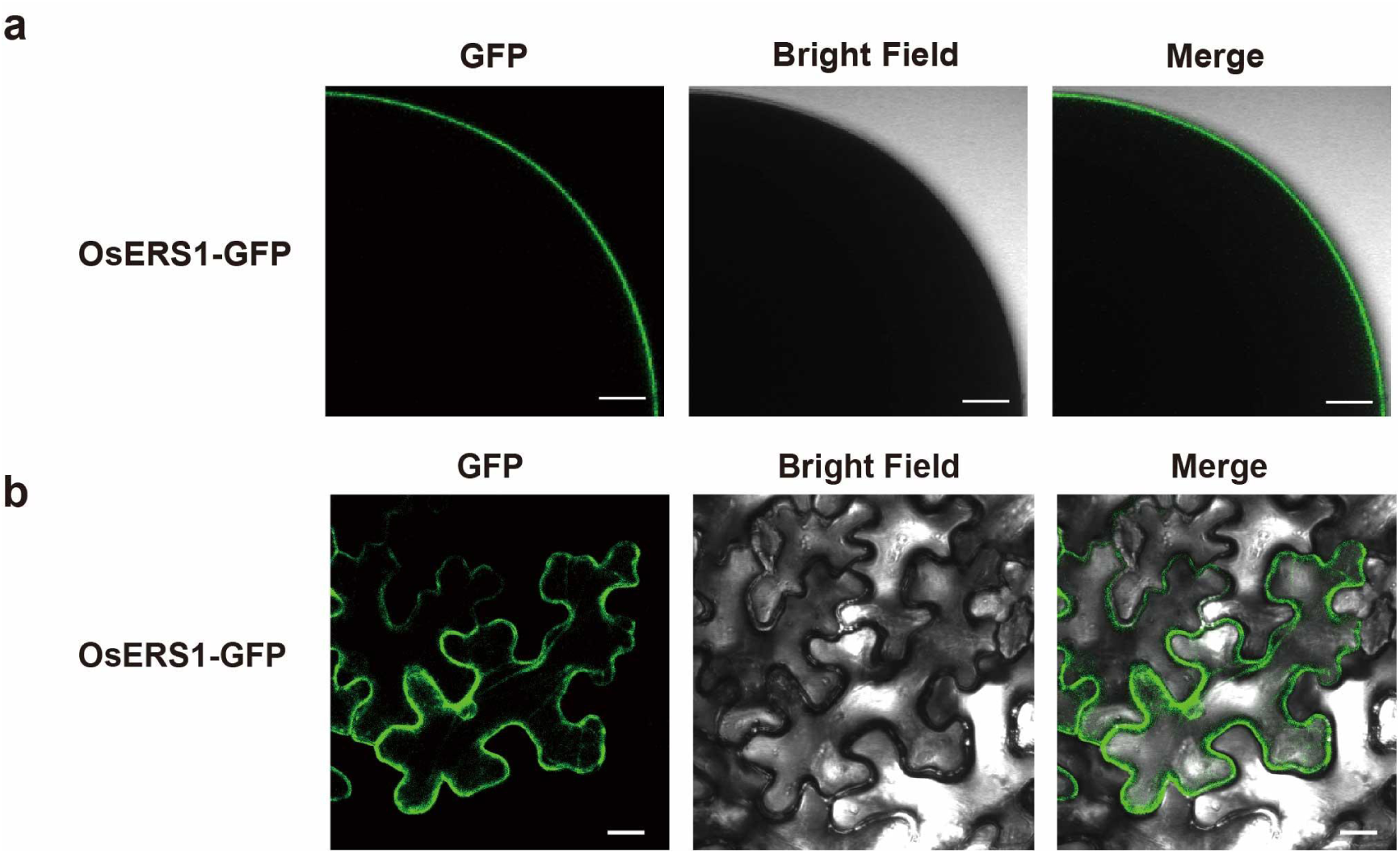
OsERS1 localizes to the plasma membrane in both *Xenopus* oocytes and *N. benthamiana* leaf cells. a, Subcellular localization of OsERS1-EGFP in *Xenopus* oocyte. Bar= 20 μm. **b,** Subcellular localization of OsERS1-EGFP in *N. benthamiana*.Bar= 200 μm.

**Supplementary Figure 6.**
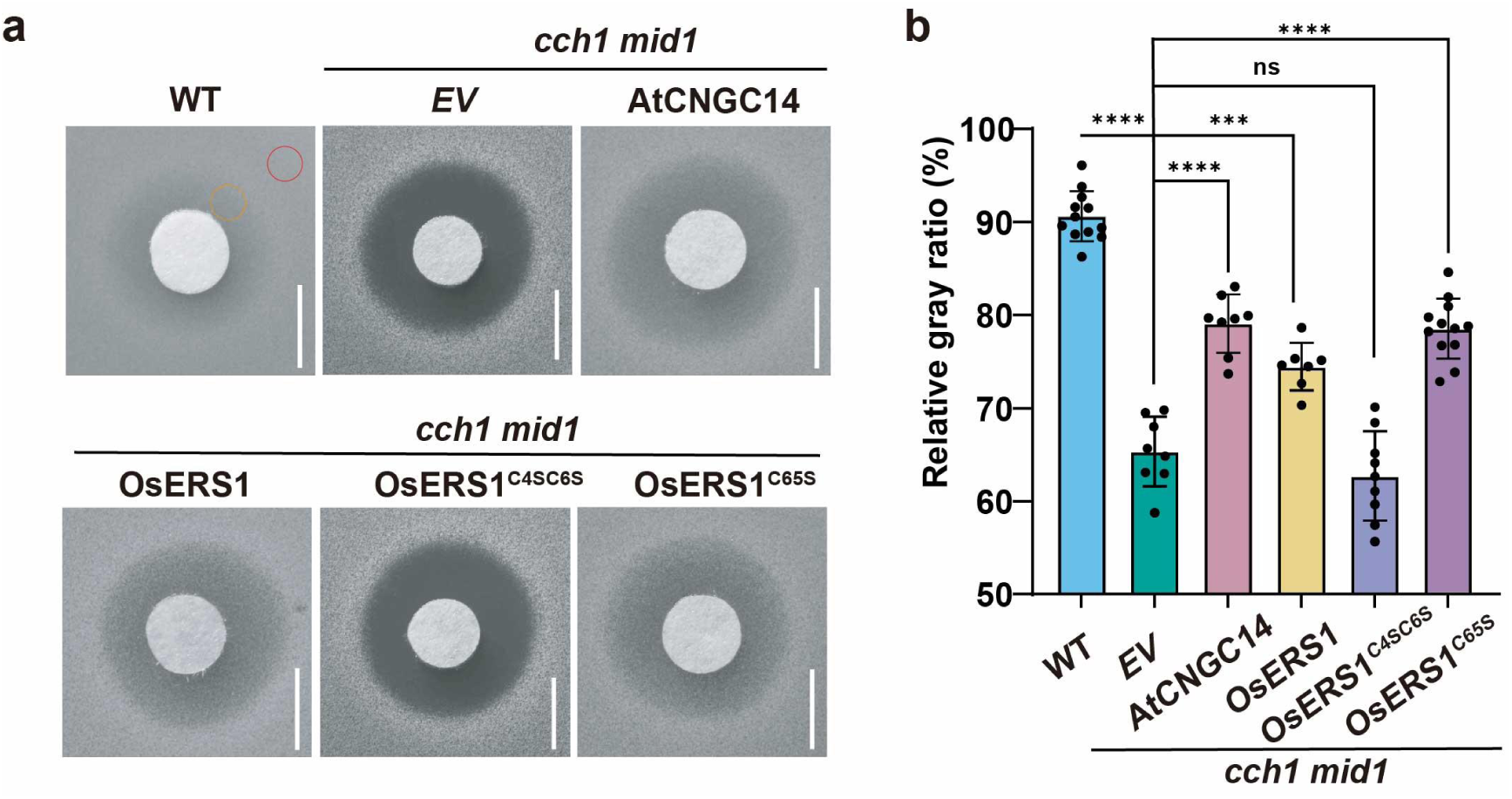
Complementation assays in calcium-uptake-deficient yeast harboring the OsERS1 point mutation. **a, b** Complementation assay in the calcium-uptake-deficient yeast mutant *cch1 mid1* transformed with empty vector (*EV*), AtCNGC14 (positive control), OsERS1, OsERS1^C4SC6S^, or OsERS1^C65S^. Data are presented as mean ± s.d. *P* values are calculated from two-sided Student’s *t*-tests. Bars = 6 mm.

**Supplementary Figure 7.**
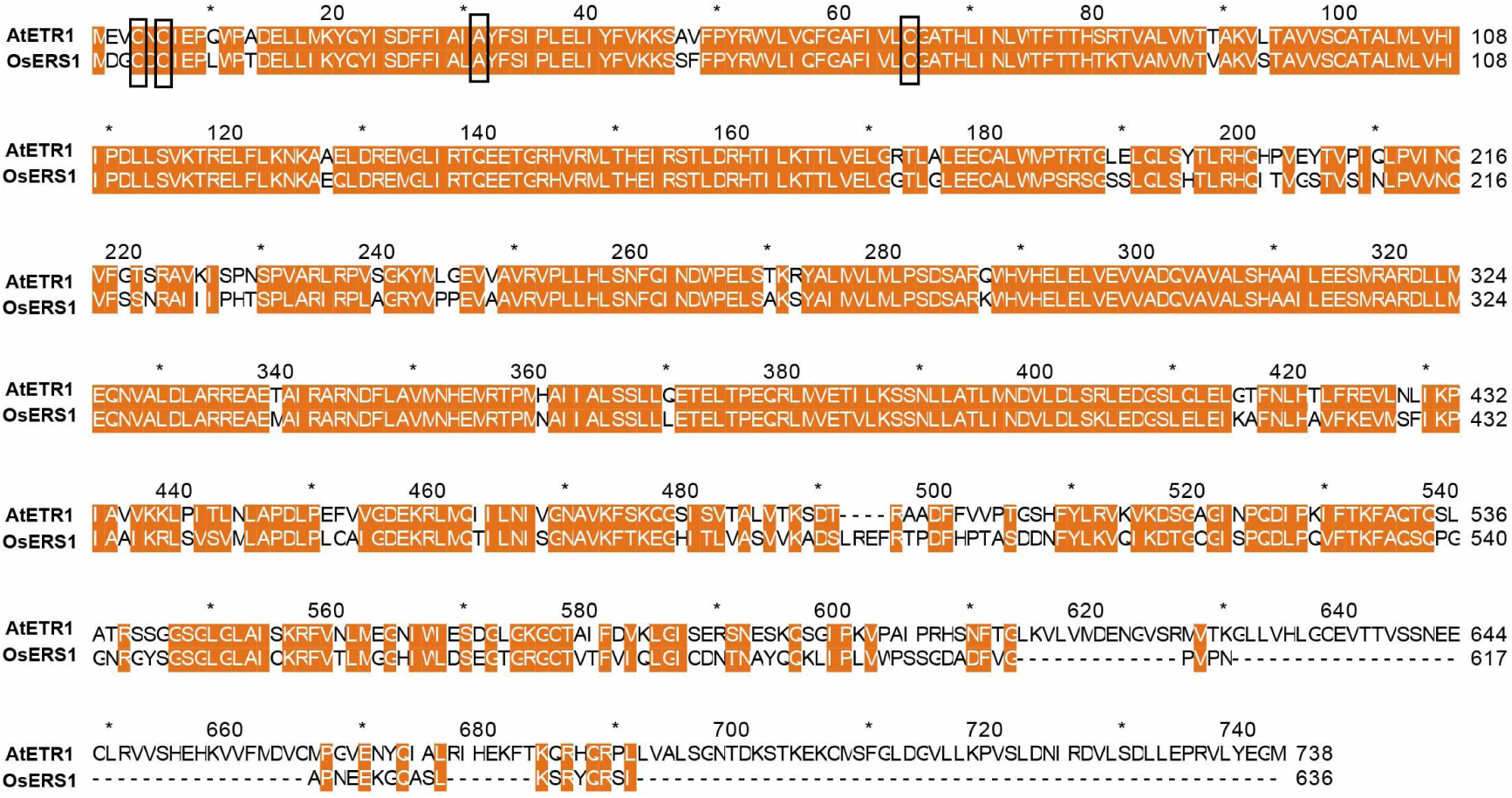
Protein sequence alignment of ethylene receptors from Arabidopsis thaliana and rice. The sequence alignment of OsERS1 and AtETR1 protein was highly conserved. The orange-colored region indicates the conserved amino acids between AtETR1 and OsERS1. The black boxes represent key residue sites, among which C4/C6 are the disulfide bond sites for dimer formation, and C65 is the key copper ion-binding site.

**Supplementary Figure 8.**
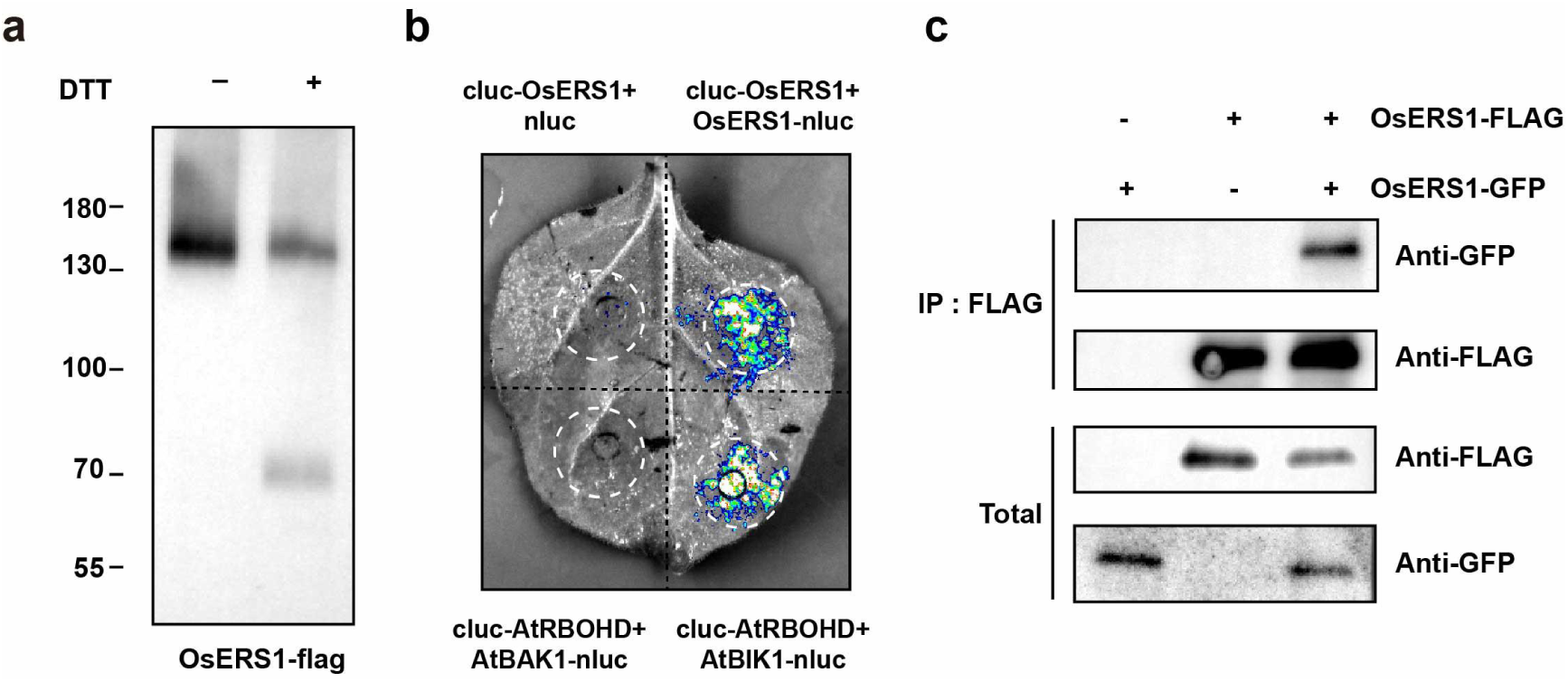
OsERS1 exhibits homomeric interaction. **a** Sensitivity of OsERS1 to reductant. Membrane fractions from *N. benthamiana* leaves were incubated in the absence (-) or presence (+) of 100 mM DTT for 1 h at 37 °C. Protein was subjected to Western Blot. **b** Split-luciferase complementation assay in *N. benthamiana* leaves confirmed OsERS1 self-interaction: Significant luciferase signals were detected in leaves co-transfected with Cluc-OsERS1 and OsERS1-Nluc. Cluc-RBOHD/BIK1-Nluc and Cluc-RBOHD/BAK1-Nluc served as positive and negative controls, respectively. **c** Co-immunoprecipitation (Co-IP) assays for interaction of OsERS1with itself. Total proteins were immunoprecipitated with Flag Trap and immunoblotted with anti-GFP and anti-Flag. IP: Immunoprecipitation.

